# Sperm competition intensity shapes divergence in both sperm morphology and reproductive genes across murine rodents

**DOI:** 10.1101/2023.08.30.555585

**Authors:** Emily E. K. Kopania, Gregg W. C. Thomas, Carl R. Hutter, Sebastian M. E. Mortimer, Colin M. Callahan, Emily Roycroft, Anang S. Achmadi, William G. Breed, Nathan L. Clark, Jacob A. Esselstyn, Kevin C. Rowe, Jeffrey M. Good

## Abstract

It remains unclear how variation in the intensity of sperm competition shapes phenotypic and molecular evolution across clades. Mice and rats in the subfamily Murinae are a rapid radiation exhibiting incredible diversity in sperm morphology and production. We combined phenotypic and genomic data to perform phylogenetic comparisons of male reproductive traits and genes across 78 murine species. We identified several shifts towards smaller relative testes mass, presumably reflecting reduced sperm competition. Several sperm traits were associated with relative testes mass, suggesting that mating system evolution selects for convergent suites of traits related to sperm competitive ability. We predicted that sperm competition would also drive more rapid molecular divergence in species with large testes. Contrary to this, we found that many spermatogenesis genes evolved more rapidly in species with smaller relative testes mass due to relaxed purifying selection. While some reproductive genes evolved rapidly under recurrent positive selection, relaxed selection played a greater role in underlying rapid evolution in small testes species. Our work demonstrates that postcopulatory sexual selection can impose strong purifying selection shaping the evolution of male reproduction, and that broad patterns of molecular evolution may help identify genes that contribute to male fertility.

## Introduction

Sperm competition often leads to the rapid evolution of male reproductive traits (Harcourt et al., 1981; Ramm et al., 2005; Pitnick et al., 2009; Lüpold et al., 2016; Pahl et al., 2018; Teves & Roldan, 2022), which can drive both extreme phenotypic divergence within populations (Pitnick et al., 2009) and the evolution of reproductive barriers between nascent species (Manier et al., 2013; Roycroft et al., 2024). Rapid evolution of reproduction also occurs at the molecular level, with reproductive genes tending to show rapid protein sequence evolution (Swanson et al., 2001; Clark & Swanson, 2005; Ahmed-Braimah et al., 2017; Dean et al., 2017; Roycroft et al., 2021a; Kopania et al., 2022). Rapid protein-coding divergence is often attributed to pervasive positive selection due to sperm competition or sexual conflict (Swanson et al., 2001; Clark & Swanson, 2005), but other processes such as relaxed purifying selection may be common for tissue-or cell-type specific male reproductive genes (Eddy, 2002; Winter et al., 2004; Larracuente et al., 2008; Larson et al., 2018; Schumacher & Herlyn, 2018; Patlar et al., 2021; Kopania et al., 2022; Murat et al., 2022). Thus, the suite of selective processes shaping the broad patterns of molecular and phenotypic evolution of male reproduction remain elusive.

If sperm competition tends to select for particular reproductive traits, then interspecific divergence in these phenotypes should be correlated with the intensity of sperm competition (Parker, 1970; Harcourt et al., 1981; Breed & Taylor, 2000; Ramm et al., 2008b; Wong, 2011; Simmons & Fitzpatrick, 2012; Lüpold et al., 2016; Varea-Sánchez et al., 2016). While directly quantifying sperm competition is challenging, many clades show increased testes mass associated with higher levels of sperm competition (Harcourt et al., 1981; Birkhead et al., 1993; Ramm et al., 2005; Firman & Simmons, 2008; Lüpold et al., 2020), thus providing a proxy for the intensity of sperm competition (Gómez Montoto et al., 2011; Pahl et al., 2018). Many studies have identified sperm morphological traits that correlate with relative testes mass, particularly in rodents [e.g., (Breed & Taylor, 2000; Immler et al., 2007; Gómez Montoto et al., 2011; Varea-Sánchez et al., 2016; Pahl et al., 2018)], but also across clades as diverse as insects and fish (Lüpold et al., 2020), suggesting that the intensity of sperm competition helps shape sperm form and function [for review see (Teves & Roldan, 2022)]. However, it remains unclear how phenotypic evolution relates to changes in molecular evolutionary rates and selective pressures across a phylogeny.

If sperm competition drives rapid divergence of reproductive proteins, then genes encoding these proteins may also evolve more rapidly with more frequent positive selection in lineages with the most sperm competition. Consistent with these predictions, molecular evolution appears to be fastest for male reproductive genes expressed during key stages of sperm morphological development (Good & Nachman, 2005; Kopania et al., 2022; Murat et al., 2022) and for secreted accessory components of the male ejaculate involved in male-female interactions and sperm competition (Dean et al., 2009). While both rapid evolution and frequent positive selection on reproductive genes is a well-established pattern, it is unclear if and how differences in the intensity of sperm competition across taxa directly correlate with lineage-specific shifts in relative evolutionary rates of reproductive genes. Do species experiencing more sperm competition and rapid phenotypic evolution also show elevated molecular evolution of reproductive genes? Observed relationships between phenotypic evolution and underlying patterns of molecular divergence between species are famously complicated (King & Wilson, 1975; Hoekstra & Coyne, 2007; Wray, 2007). Even when positive directional selection is pervasive on a given trait, we expect that selection of the underlying genes will be relatively rare genome-wide (Hernandez et al., 2011; Harris et al., 2018). Given this, variation in purifying selection is expected to be the primary determinant of genome-wide patterns of molecular evolution (Kimura, 1968; Eyre-Walker & Keightley, 2007; Larracuente et al., 2008). However, it is unclear how rapid trait evolution driven by positive selection changes the overall landscape of purifying selection on non-positively selected genes that are functionally associated with that trait. For reproductive genes, there is strong evidence for positive directional selection on specific functional groups of genes (Clark et al., 2006), including those involved in fertilization (Swanson et al., 2001), sperm swimming and flagellar movement (Podlaha et al., 2005), and seminal fluid (Clark & Swanson, 2005; Good et al., 2013). Some positively selected reproductive genes show evidence for elevated rates of positive selection in lineages with higher levels of sperm competition (Dorus et al., 2004; Ramm et al., 2008b; Finn & Civetta, 2010; Wong, 2010, 2014), but others do not show this pattern (Ramm et al., 2008b; Finn & Civetta, 2010; Good et al., 2013; Carnahan-Craig & Jensen-Seaman, 2014), and therefore it is not known if this pattern is common across positively selected reproductive genes (Wong, 2011). Additionally, recent studies have shown that relaxed purifying selection can play an important role in the rapid evolution of reproductive genes (Dapper & Wade, 2020; Johnson et al., 2022; Bowman et al., 2024; Zhao et al., 2024), but it is unclear how changes in sperm competition intensity across lineages shapes the molecular evolution of different functional classes of reproductive genes. Is positive selection on traits associated with changes in mating system generally accompanied by enhanced or relaxed purifying selection on the genes that underlie the development of that trait?

The subfamily Murinae represents one of the most rapid mammalian radiations with >700 species (i.e., >10% of extant mammal species) arising over the last ∼12-14 million years (Rowe et al., 2016; Rowe et al., 2019; Roycroft et al., 2021a). Murines also show striking diversity in reproductive biology (Breed et al., 2020b; Roycroft et al., 2021a), including extreme interspecific divergence in relative testes mass and sperm morphology [Figure 1; (Breed, 2005)]. The reasons for this diversity of reproductive traits are not fully understood, but there is some evidence that murine species with shorter mating seasons and less spatial clustering of females tend to have males with larger relative testes mass (Firman et al., 2022). The house mouse (*Mus musculus*) and Norway rat (*Rattus norvegicus*) model species are part of this radiation, providing extensive genomic resources (Gibbs et al., 2004; Keane et al., 2011; Roycroft et al., 2021a) and a detailed understanding of murine spermatogenesis and reproductive biology (Green et al., 2018; Breed et al., 2020b; Firman, 2020). Previous studies suggest that some sperm morphological traits have evolved in response to sperm competition in murine rodents (Gómez Montoto et al., 2011; Varea-Sánchez et al., 2016; Pahl et al., 2018), but these works relied on phylogenies that included relatively few species, were not well-resolved, or were inferred based on a small number of genomic regions (Fabre et al., 2012; Steppan & Schenk, 2017). Thus, comparative analyses of murine rodent reproductive evolution have been limited in their ability to connect phenotypic and genome-wide molecular evolution with changes in sperm competition levels across many species. Here, we use exome sequencing across 33 murine species, combined with published exome (Sarver et al., 2017; Roycroft et al., 2021a; Roycroft et al., 2021b) and genome (Gibbs et al., 2004) data for 45 species, resulting in 11,775 sequenced protein-coding genes for 78 species with available reproductive trait data. We address four main questions: (i) How does the intensity of sperm competition shape variation in sperm morphology in murine rodents? (ii) Is there heterogeneity of molecular evolutionary rates among reproductive tissues and cell types across murines? (iii) What are the relative roles of positive selection and relaxed purifying selection in driving the molecular evolution of male reproductive genes? (iv) Are changes in the intensity of sperm competition associated with lineage-specific shifts in molecular evolutionary rates?

**Figure 1.**
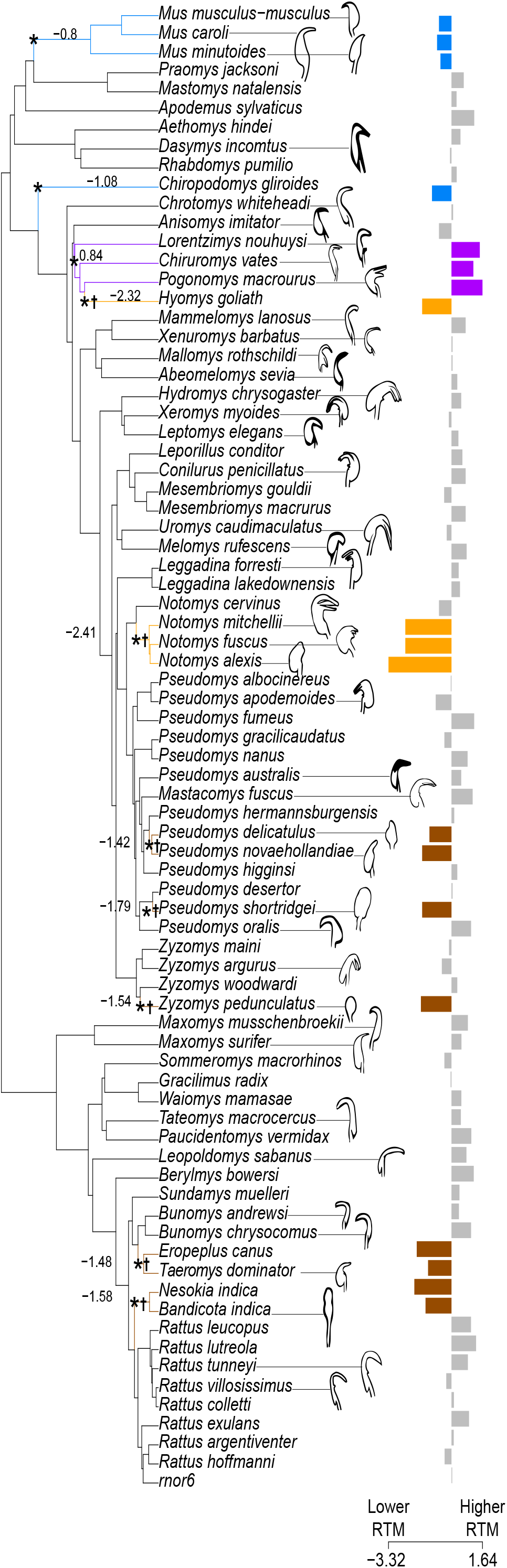
Phylogenetic variation in relative testes mass and sperm head morphology among 78 murine species. Bar plots show normalized values of relative testes mass, with different colors representing distinct adaptive peaks based on an OU model. Blue, orange, and brown bars indicate shifts to smaller testes, while purple bars indicate shifts to larger testes. (*) indicate phylogenetic nodes corresponding to shifts in phenotypic optima, and adjacent values show optimum trait values at these nodes. (†) show lineages used as foreground small testes species for downstream analyses. Traces of sperm micrographs (not to scale) are shown for some species to highlight the diversity of sperm head morphology.

## Materials and Methods

### Samples and data collection

This study was part of a more extensive phylogenomics analysis of 210 murine species. Here we focus on the subset of these species with sufficient reproductive data, and analysis of the full dataset will be reported elsewhere. For the specimens included in this study, we combined published data with new data collected from museum specimens for a total of 78 species representing all the major tribes within Murinae (accession numbers and museum catalogs in Supplemental Table S1). All specimens collected after 2012 have material transfer agreements and are compliant with the Nagoya Protocol on Access and Benefit Sharing. Over 80% of samples were sourced from Australia (45%) or Indonesia (40%) as part of a long-term collaborative partnership with authors from those countries and who are in involved in this study (ASA; KCR). We extracted DNA using a Qiagen DNeasy kit with modifications to account for tissue preservation [Supplemental Methods; (Shapiro & Hofreiter, 2012; Bi et al., 2013)]. We performed library preparation using a Kapa Biosystems HyperPrep Kit (Roche Diagnostics Corporation, Indianapolis, IN) and exome capture using custom SeqCap EZ Developer Probes (Roche Diagnostics Corporation, Indianapolis, IN) targeting 203,188 exon regions (mm9 annotation). Pooled libraries were sequenced on Illumina NextSeq at HudsonAlpha Institute for Biotechnology, HiSeq 4000 at the University of Oregon, and HiSeqX at Novogene. For Australian samples, libraries were prepared using a modified Meyer & Kircher protocol (Meyer & Kircher, 2010), following the methods described in (Roycroft et al., 2019; Roycroft et al., 2021b) and sequenced on an Illumina HiSeq 2500 at the Australian National University’s Biomolecular Resource Facility. We used liftOver to transfer the coordinates for targeted regions to match the mm10 reference genome (Hinrichs et al., 2006). We also used data from 44 published exomes (Sarver et al., 2017; Roycroft et al., 2021a; Roycroft et al., 2021b) and the *Rattus norvegicus* reference assembly [rnor6; (Gibbs et al., 2004)]. Phenotypic data were compiled from several publications (Supplemental Table S2; Supplemental Methods).

### Assembly, alignment, and phylogenetic inference

Because this work was part of a more extensive phylogenomic analysis of murine rodents, we assembled whole exomes using our full 210 species dataset, performed alignment and phylogenetic inference on a set of 188 species that passed filtering, and then pruned this 188 species tree to include only the 78 species for which we have reproductive phenotype data. We first removed adaptors and removed low-quality reads using the program fastp (Chen et al., 2018). We performed *de novo* exome assembly on samples from 210 species using SPAdes (Bankevich et al., 2012) and assessed assembly quality and corrected low-quality bases using *Referee* (Thomas & Hahn, 2019). We re-mapped reads to the corrected assemblies to genotype each sample. Exome assemblies were annotated using transcripts from mouse (mm10) and rat (rnor6) reference genomes that exist in both references as one-to-one orthologs with an Ensembl orthology confidence of 1 and have an *d*_S_ value <0.5 between mouse and rat. When multiple transcripts passed the first two filters, we kept the one containing the most probe targets. Transcripts representing about 65% of targeted genes passed filtering.

To generate alignments, we first identified homologous regions between mouse reference exons and our assembled contigs using BLAST. We then converted mouse reference exons to trimmed amino acid sequences to keep coding sequences between exons in-frame, and we used exonerate (Slater & Birney, 2005) to identify exons in our assembled contigs that were homologous to mouse reference exons. We filtered out exons with fewer than 175 samples with a matching BLAST hit and aligned the remaining exons using MAFFT (Katoh et al., 2002). We then back-translated to nucleotide sequences and filtered out sequences that were >20% gaps or had alignment sites in 3-codon windows in which 2 or more codons had 2 or more gaps in over half of the sequences. Lastly, we filtered out samples that had premature stop-codons after alignment filtering. After filtering, our alignments had on average > 150 aligned sequences per protein, and the average non-gapped sequence length per protein was about 250 codons.

After extensive filtering, our experiment recovered multispecies alignments for 11,775 of 18,283 (64.5%) protein-coding genes targeted by the mouse whole exome capture design. To provide an empirical assessment of overall alignment qualities, we estimated *d*_S_ for each gene using the HyPhy MG94 fit model. We found that the overall distribution of *d*_S_ values was not strongly skewed and that all genes included in our post-filtering datasets had *d*_S_ values < 0.05, consistent with generally high-quality alignments in the post-filtered gene set. Another important consideration for analyses of molecular evolution is how the experimental details of exome capture may bias resulting genic patterns of molecular evolution. Most of the filtered genes were excluded *a priori* based on annotated uncertainty in orthology between mouse and rat (82.4%, 5340 of 6481 filtered genes based on available annotation), and not because they showed poor capture recovery across species. Only ∼6% of genes were filtered based on incomplete recovery or details of their molecular evolution (e.g., elevated substitution rates, stop codons), and these estimates were not highly variable or obviously biased across reproductive tissues [range 3.5% (prostate) to 9% (somatic testis expression); (Supplemental Table S3)]. Although capture performance could theoretically be sensitive to extensive sequence evolution, probe-based hybridization and enrichment is generally expected to be robust to the moderate levels of sequence divergence observed between murine species (Jones & Good, 2016).

Our post-filtering dataset included 188 species and 11,775 protein-coding genes. From these alignments, we inferred gene trees using IQ-TREE v2.0.4 (Nguyen et al., 2014; Kalyaanamoorthy et al., 2017) with 1000 bootstrap iterations (Hoang et al., 2017) and inferred a summary species tree from the gene trees using ASTRAL-MP v5.15.2 (Yin et al., 2019). We estimated branch lengths in terms of relative number of substitutions on the ASTRAL species tree using loci whose inferred gene tree had a normalized Robinson-Foulds distance to the species tree of less than 0.25 to minimize the effects of incomplete lineage sorting on branch length estimation (Mendes & Hahn, 2016). We then pruned this summary species tree using *ape* (Paradis & Schliep, 2018) to generate a species tree with 78 taxa that had male reproductive phenotype data (Figure 1).

### Phylogenetic analyses of trait evolution

We modeled the evolution of relative testes mass using an OU model implemented in *l1ou* v1.43 and identified convergent shifts in phenotypic optima using the function *estimate_convergent_regimes* with *criterion*=“AIC” and *method*=“rr” (Khabbazian et al., 2016). We tested if sperm morphology parameters were significantly associated with relative testes mass using phylogenetic logistic regression (Ives & Garland, 2009) in *phylolm* v2.6.2 (Tung Ho & Ané, 2014) for binary sperm hook presence/absence or phylogenetic generalized least squares in *nlme* v3.1.160 (Pinheiro J, 2020) for all other traits. We evaluated three different correlation structures: Brownian motion, Pagel’s ƛ (Pagel, 1999), and an OU model (Martins & Hansen, 1997). We did not have all phenotype data for all species, so we pruned the tree for each trait to include only taxa with phenotype data for that trait (Supplemental Table S4). We used R v4.1.1 for all trait evolution analyses.

### Phylogeny-wide molecular evolution

We calculated rates of protein sequence evolution (*d*_N_/*d*_S_) and tested for gene-wide positive selection across the entire phylogeny using the BUSTED (Murrell et al., 2015) model in HyPhy v2.5.42 (Kosakovsky Pond et al., 2019) without specifying any foreground branches. We input alignments and unrooted gene trees that included species with reproductive phenotype data (Figure 1). To generate these trees, we pruned individual gene trees to include only species in the 78-species reproductive phenotype dataset using the *tree_doctor --prune* function in PHAST (Hubisz et al., 2010) and unrooted trees using Newick Utilities (Junier & Zdobnov, 2010). To improve runtimes, we removed duplicate sequences when present using the HyPhy “remove-duplicates” script (https://github.com/veg/hyphy-analyses/tree/master/remove-duplicates). Tests for positive selection may be prone to false positives due to synonymous rate variation across sites within a gene and multiple-nucleotide substitutions, which are often excluded from substitution models (Wisotsky et al., 2020; Lucaci et al., 2023). However, models that incorporate these parameters may be a worse fit for the data for some loci, and introducing additional parameters can potentially reduce power (Lucaci et al., 2023). Therefore, we ran BUSTED three times: (1) with the settings “--srv Yes” and “--multiple-hits Double+Triple” to model synonymous rate variation across sites and multiple nucleotide mutations, (2) with the settings “--srv Yes” and “--multiple-hits None” to model synonymous rate variation only, and (3) with the settings “--srv No” and “--multiple-hits None” to model neither. We then calculated the model averaged p-value across all three models, or the sum of p-values weighted by their Akaike Information Criterion score. This method incorporates model likelihood into p-value calculation and has been shown to reduce false positives while having similar power to models that do not include synonymous rate variation or multiple-nucleotide substitutions (Lucaci et al., 2023). We filtered out loci for which the models did not converge (likelihood ratio test < 0), corrected the remaining model-averaged p-values for multiple tests using Benjamini-Hochberg correction, and considered genes with adjusted model-averaged p-values < 0.05 to be under positive selection. Gene-wide estimates of *d*_N_/*d*_S_ were similar across all three BUSTED models, so we report the *d*_N_/*d*_S_ values estimated by the “--srv Yes” and “--multiple-hits Double+Triple” model. To ensure that our inferences of rapid evolution were not being driven by genes with high dS values, which could indicate lower quality alignments, we identified genes with the highest 2% dS values (dS > 0.034). Among these genes, the maximum *d*_N_/*d*_S_ value was 1 and the distribution of *d*_N_/*d*_S_ values was similar to that of all genes, suggesting that genes with high *d*_N_/*d*_S_ values in our dataset are not the result of low-quality alignments.

We tested if molecular evolutionary rates varied across accessory male reproductive tissues by calculating median *d*_N_/*d*_S_ values for genes enriched in these tissues. We considered a gene enriched in a tissue if its protein product was detected at the highest level in that tissue relative to other male reproductive tissues based on *M. musculus* protein mass spectrometry data reported by (Dean et al., 2009). Testes were not included in this proteomics dataset, so we used a set of testis-specific genes from (Chalmel et al., 2007), which identified genes expressed in testis but not in a set of 17 somatic tissues, cumulus-oocyte complex cells, or ovaries. To compare evolutionary patterns in testis-specific genes to those of other tissue-specific genes, we used a list of *M. musculus* tissue-specific genes (Li et al., 2017). We also compared *d*_N_/*d*_S_ for genes enriched for expression across different testes cell types using *M. musculus* single-cell RNAseq data from (Green et al., 2018). Genes were considered enriched in a cell type if they showed >20% detection rate, >2-fold mean expression levels, and a significant binomial likelihood test when comparing their expression in a focal cell type to their expression in all other testes cell types. See Supplemental Methods for details on tissue and cell-type gene sets. We then calculated median *d*_N_/*d*_S_ values across all genes in each of these tissue-or cell type-associated gene sets and compared these values to the genome-wide *d*_N_/*d*_S_, calculated as the median *d*_N_/*d*_S_ across all genes in our dataset, using a Wilcoxon rank sum test with Benjamini-Hochberg correction for multiple tests. We then compared the proportions of genes with evidence for selection for these gene sets to the genome-wide average. For spermatogenesis cell types, we also calculated the proportion of genes enriched in each cell type that were considered testis-specific in (Chalmel et al., 2007). From these lists of genes associated with reproductive tissues or cell types, the proportions of genes that passed filtering were largely consistent with the genome-wide pass rate of 65%. However, some groups had lower pass rates because a higher proportion of genes lacked a confident one-to-one ortholog between mouse and rat, likely reflecting the high paralogy of some reproductive genes (Supplemental Table S3).

### Lineage-specific molecular evolution

We next used *RERconverge* (Kowalczyk et al., 2019) to identify genes showing relative molecular evolutionary rate shifts across lineages associated with convergent transitions in relative testes mass. Because our OU models primarily identified convergent adaptive shifts to lower testes mass, we set species consistently identified as having significant shifts to smaller testes as foreground species (Figure 1). We also ran *RERconverge* with large testes mass species (top 10% of relative testes mass values) as foreground species and with relative testes mass as a continuous trait. We verified results from *RERconverge* using 1000 permutations of phylogenetic simulations [i.e., permulations; (Saputra et al., 2021)] with a Benjamini-Hochberg correction for multiple tests. We then used *topGO* (Alexa et al., 2006) to test for gene ontology (GO) term enrichment among the top 250 most accelerated genes. To visualize enrichment for the spermatogenesis GO term among the top most accelerated genes, we ranked genes by their *RERconverge* Rho values and plotted the tricube moving average of enrichment for spermatogenesis genes in sliding windows along these ranked genes. The tricube moving average is a smoothing function that down-weights values at the ends of each window to limit the effects of extreme values (Kowalczyk et al., 2020).

Accelerated evolutionary rates associated with a convergent phenotype could be due to positive selection or relaxed purifying selection in lineages with the phenotype. Therefore, we tested for episodic positive selection or relaxed purifying selection on spermatogenesis genes in small testes species with the BUSTED-PH (https://github.com/veg/hyphy-analyses/tree/master/BUSTED-PH) and RELAX (Wertheim et al., 2014) models in HyPhy, with small testes mass species as “test” branches and all other species as “background”. Internal branches for which all descendent species were in the small relative testes mass group were also considered test branches, and all other internal branches were considered background. BUSTED-PH explicitly tests if episodic positive selection is associated with test branches, but not background branches, to distinguish convergent positive selection from phylogeny-wide positive selection (Kowalczyk et al., 2021). We ran both BUSTED-PH and RELAX using the same pruned, unrooted gene trees we used to run BUSTED. We also ran these tests with synonymous rate variation and multiple-nucleotide mutations incorporated into the models. Genes with BUST-ED-PH p-values < 0.05 after Benjamini-Hochberg correction were considered to be under episodic positive selection in species with small relative testes mass. Genes with RELAX p-values < 0.05 after Benjamini-Hochberg correction and < 1 were considered to be under relaxed purifying selection, and genes with RELAX corrected p-values < 0.05 and K > 1 were considered to be under intensified selection on test branches. Two genes that had significant p-values under both the BUSTED-PH and RELAX models were placed in the “positive selection in small testes mass species” category.

## Results

### High quality assemblies and phylogeny of diverse murine rodents

Our post-filtering dataset included 188 murine species and 11,775 protein-coding genes, which we used to generate assemblies, alignments, gene trees, and a species tree that will be reported in a forthcoming manuscript. Here, we report results for 78 species with both exome (n=77) or genome (n=1) sequences and male reproductive phenotypes plus two outgroups (Supplemental Table S1, Supplemental Table S2). Across the 77 exomes, about 89% of reads on average mapped to our *de novo* assemblies with an average depth of 22×. Based on *Referee* (Thomas & Hahn, 2019), our assemblies had on average fewer than 0.0005 errors per base and about 2% low quality positions, indicating that our assemblies were of high quality. The resulting species tree (Figure 1) was well-supported, with one node having an ASTRAL support value of 0.93 and all other nodes having an ASTRAL support value of 1. However, concordance factors were more varied, with a median gene concordance factor of 38 and a median site concordance factor of 45.7, likely reflecting high levels of incomplete lineage sorting or introgression (Hibbins et al., 2020).

### Murine rodents show repeated shifts to smaller relative testes mass

Our findings suggest considerable variation in mating behavior and likely sperm competition (Breed & Taylor, 2000), with several species groups independently evolving mating systems with lower levels of multiple mating. Relative testes mass ranged from 0.1% to 4.8% of body mass (Figure 1; Supplemental Table S2). The Ornstein-Uhlenbeck (OU) model identified four adaptive peaks corresponding to convergent phenotypic optima. Three peaks represented shifts to smaller relative testes mass (blue, orange, and brown; Figure 1) and one represented a shift to larger relative testes mass (purple; Figure 1), including multiple examples of convergence on the same optimum (i.e., the same peaks on independent branches). Because three of the four shifts were to smaller testes mass, we focused on small testes as our foreground trait. OU models can be prone to errors in estimates of adaptive peaks (Cooper et al., 2016), and we found that the number of adaptive peaks for species with small testes mass was sensitive to model selection (Supplemental Figure S1). Indeed, one of these adaptive shifts (blue, Figure 1) had inconsistent species membership across different model selection methods (Supplemental Figure S1), represented a small phenotypic optimum, and included *Mus musculus*, which experiences considerable sperm competition in natural populations (Dean et al., 2006). Given these considerations, we focused on the two groups with the most pronounced phenotypic shifts (i.e., orange and brown groups denoted by † in Figure 1) as representing murines with the lowest relative levels of sperm competition. These groups were combined as one “low sperm competition” group for further analyses.

### Sperm competition selects for longer sperm hooks, narrower heads, and longer tails

We used phylogenetic logistic regression [binary traits; (Ives & Garland, 2009)] or phylogenetic generalized least squares (Martins & Hansen, 1997) to test if sperm morphology traits were correlated with relative testes mass. We found that hook presence was positively associated with relative testes mass (FDR-corrected P < 0.01; Figure 2A). We ran phylogenetic generalized least squares using Brownian motion, Pagel’s λ (Pagel, 1999), and an OU model (Martins & Hansen, 1997). With all three models, we found significant associations between relative testes mass and apical hook length, apical hook angle, ventral process length, ventral process angle, head width, tail length (i.e., flagellum length), and principal and end piece length (FDR-corrected P < 0.05; Figure 2; Supplemental Figure S2; Supplemental Table S4). Sperm head width was negatively associated with relative testes mass (FDR-corrected P < 0.01; Figure 2E), suggesting that sperm competition selects for narrower sperm heads with faster swimming speed (Hook et al., 2021). Sperm head length and sperm head area were correlated with relative testes mass under some models (Supplemental Figure S2; Supplemental Table S4). Some of these sperm morphological traits are correlated with each other, likely due to size scaling relationships (Supplemental Table S5), and therefore may not represent independent targets of selection. Collectively, these data reveal correlated evolution across traits that are likely relevant to sperm competitive ability and, to the extent that relative testes mass is a robust proxy of sperm competition, reflect phenotypic evolution in response to lineage-specific variation in the intensity of sperm competition.

**Figure 2.**
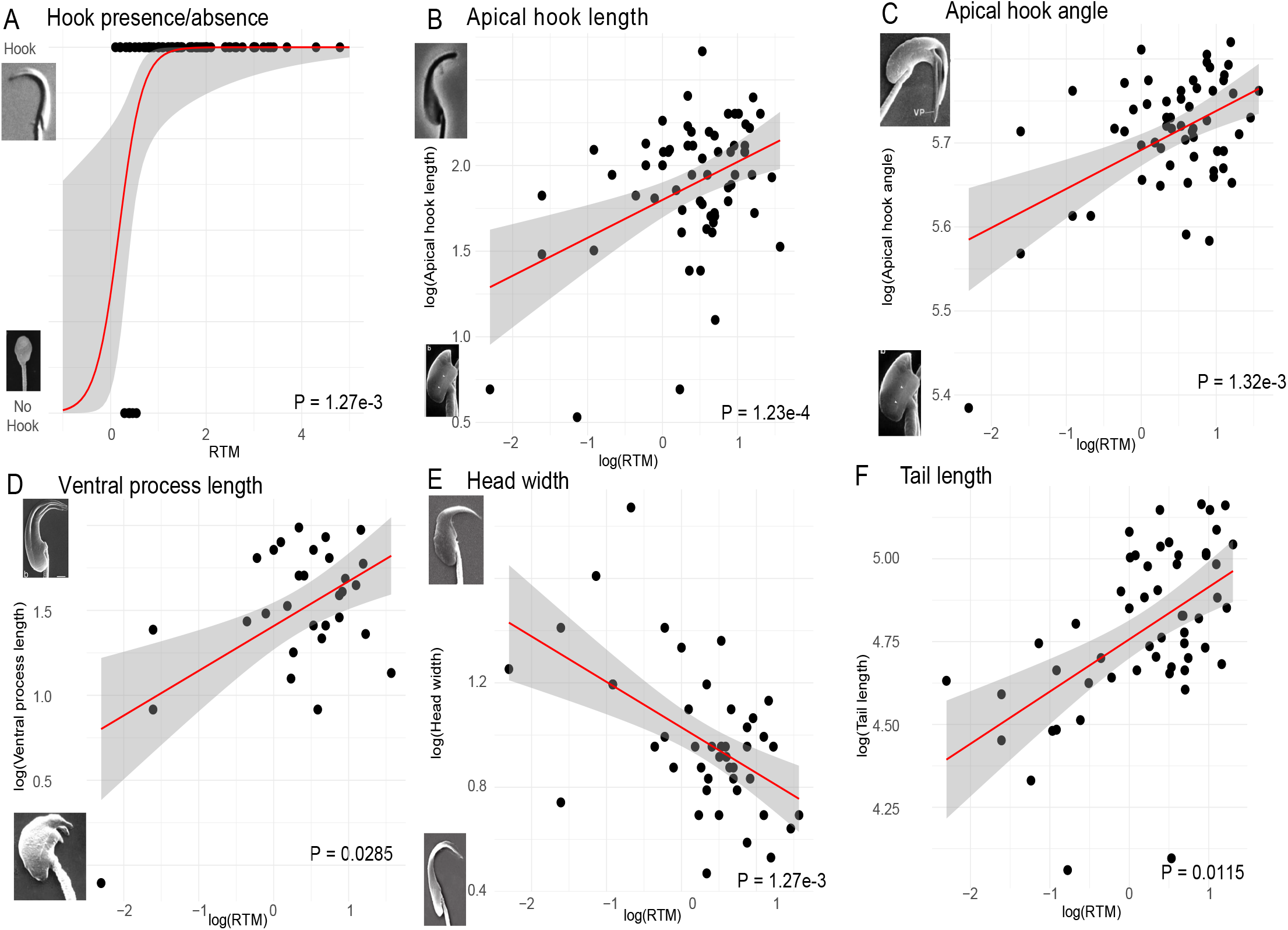
Correlations between relative testes mass (RTM) and sperm morphological traits. Red lines and gray areas show regressions and confidence intervals based on a logistic regression (A) or generalized linear models (B-F). P-values are based on phylogenetic logistic regression (A) or phylogenetic generalized least squares analyses using a Brownian motion model (B-F) with Benjamini-Hochberg correction for multiple tests. All lengths and widths were measured in μm, angles were measured in degrees, and relative testes mass was measured as a percent of body mass. Sperm micrographs show examples of the extreme ends of each trait along the y-axis (sources in Supplemental Table S4).

### Rapid evolution of genes enriched in late spermatogenesis and seminal vesicles

Next, we evaluated molecular evolution across different components of male reproduction. We calculated rates of protein-coding evolution (*d*_N_/*d*_S_) for genes grouped based on the male accessory tissue (seminal vesicles, coagulating gland, dorsolateral prostate, ventral prostate, bulbourethral gland, bulbourethral diverticulum) in which their encoded proteins were detected at the highest abundance in proteomics data (Dean et al., 2009). We also estimated an average *d*_N_/*d*_S_ for genes with testis-specific expression patterns in house mice (Chalmel et al., 2007). Most male reproductive tissues had average *d*_N_/*d*_S_ values similar to or lower than the genome-wide average (Table 1; Figure 3A), suggesting protein sequence conservation (Dean et al., 2009). However, seminal vesicle genes tended to evolve rapidly (*d*_N_/*d*_S_ = 0.29; Table 1; Figure 3A; FDR-corrected Wilcoxon rank-sum P < 0.05). The median *d*_N_/*d*_S_ of all genes detected in the seminal vesicles was not significantly higher than the genome-wide average (Supplemental Table S6), suggesting that the more rapid divergence was limited to genes specialized to this tissue. Testis-specific genes had an average *d*_N_/*d*_S_ of 0.19, higher than the genome-wide average (Figure 3A; FDR-corrected Wilcoxon rank-sum test P < 0.05). Testis-specific genes may evolve rapidly because they are tissue-specific and therefore released from pleiotropic constraint. To test this, we compared the median *d*_N_/ *d*_S_ of testis-specific genes to those of other tissue-specific genes (Li et al., 2017) and found that testis-specific genes had significantly higher *d*_N_/*d*_S_ than ovary-specific genes or a combined set of all tissue-specific genes (Supplemental Table S7). Thus, testis-specific genes tended to evolve even more rapidly than other tissue-specific genes. Seminal vesicle genes and testis-specific genes both had relatively high proportions of genes filtered out due to lack of 1:1 orthology between mouse and rat (54% and 41%, respectively, compared to a genome-wide average of 29%; Supplemental Table S3). This suggests that the copy numbers of these genes may be evolving rapidly, in addition to their protein sequence divergence.

**Table 1.** Summary of molecular evolution by spermatogenesis stage and male reproductive tissue. (*) indicate significant differences from the average of all genes (FDR-corrected P < 0.05). SO = somatic; SG = spermatogonia; PL = pre-leptotene; SC = spermatocytes; SD = spermatids; EL = elongating; BD = bulbourethral diverticulum; BG = bulbourethral gland; CG = coagulating gland; DP = dorsolateral prostate; VP = ventral prostate; SV = seminal vesicle; TS = testis-specific; _a_ (Green et al., 2018); _b_ (Dean et al., 2009); _c_ (Chalmel et al., 2007)

|  | All Genes | Spermatogenesis cell types<sup>a</sup> |  |  |  |  |  | Male reproductive tissues |  |  |  |  |  |  |
| --- | --- | --- | --- | --- | --- | --- | --- | --- | --- | --- | --- | --- | --- | --- |
|  | All Genes | SO | SG | PL | SC | SD | EL | BD<sup>b</sup> | BG<sup>b</sup> | CG<sup>b</sup> | DP<sup>b</sup> | VP<sup>b</sup> | SV<sup>b</sup> | TS<sup>c</sup> |
| Number of genes | 10610 | 57 | 438 | 227 | 95 | 228 | 186 | 43 | 131 | 1 | 72 | 36 | 7 | 378 |
| Median $d_N/d_S$ | 0.12 | 0.07* | 0.09* | 0.10* | 0.13 | 0.19* | 0.16* | 0.08 | 0.10* | 0.09 | 0.09 | 0.13 | 0.29* | 0.19* |
| Percent of genes under positive selection | 17% | 18% | 17% | 19% | 17% | 14% | 16% | 19% | 18% | 0% | b% | 14% | 14% | 13% |
| Percent of testis-specific genes under positive selection | 13% | 0% | 0% | 5.6% | 9.1% | 12% | 17% | NA | NA | NA | NA | NA | NA | NA |
| Percent of genes that are testis-specific | 3.6% | 3.5% | 1.4%* | 7.9%* | 23%* | 32%* | 25%* | NA | NA | NA | NA | NA | NA | NA |

**Figure 3.**
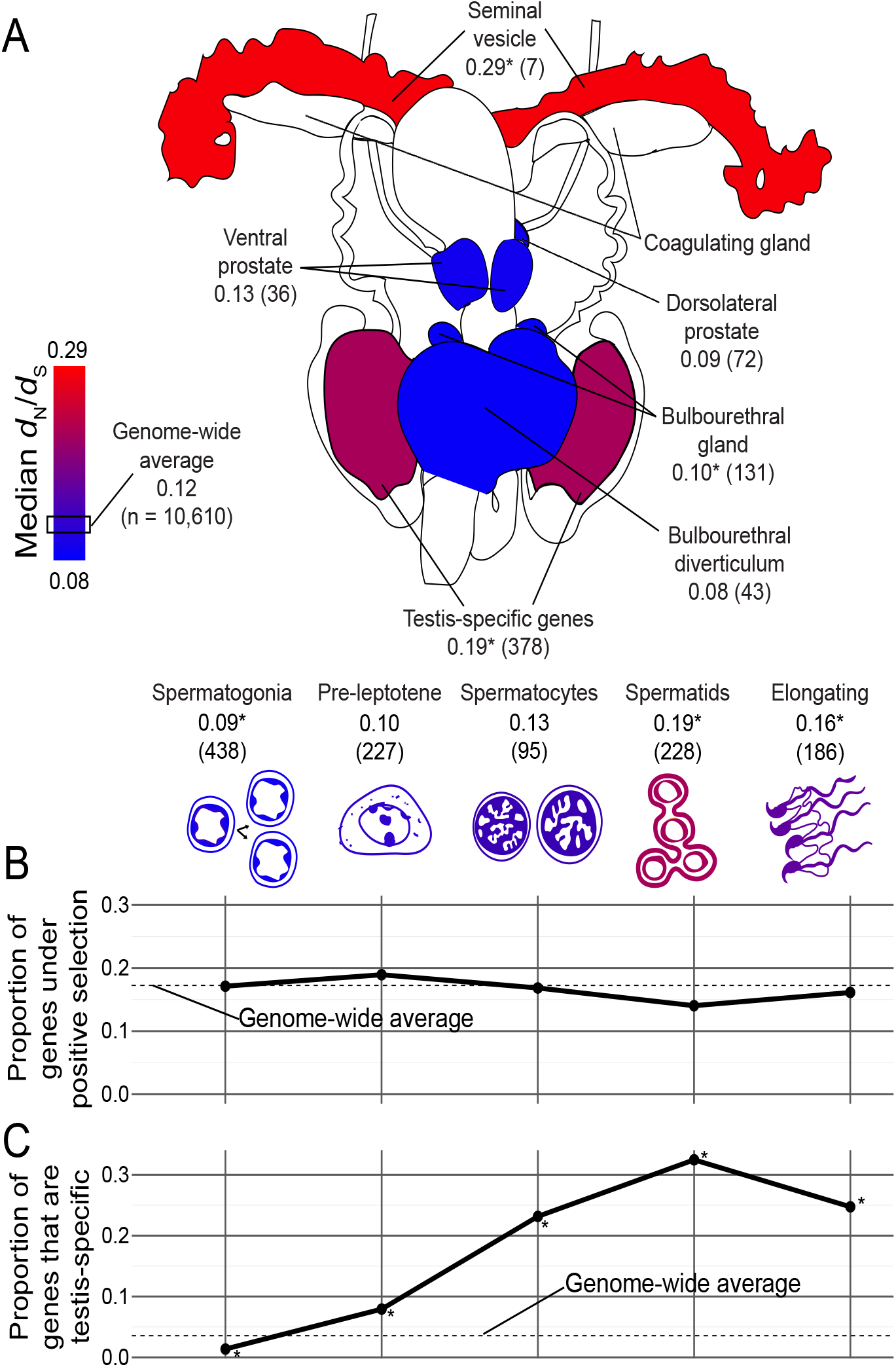
Molecular evolution by male reproductive tissue and spermatogenesis stage. (A) Mouse male reproductive tract and spermatogenesis cell types ordered from earliest to latest developmental stages. Numbers represent median *d*_N_/*d*_S_ values for each tissue or cell type with gene counts in parentheses. Coagulating gland was excluded because data were only available for one gene in this tissue. The box on the scale indicates the median phylogenywide *d*_N_/*d*_S_ (0.12) across all genes, and (*) indicate significant differences from this median *d*_N_/*d*_S_ (Wilcoxon rank sum test FDR-corrected P < 0.05). (B) Proportion of genes enriched in each cell type with evidence for positive selection based on BUSTED model-averaged p-values. The dotted line represents the proportion of genes with evidence for positive selection out of all genes included in the test for selection (no cell types were significantly different from the genome-wide average; FDR-corrected Pearson’s chi-squared P > 0.05). (C) Proportion of genes enriched in each cell type that were testis-specific, with (*) indicating significance as above. Figure concept and male reproductive tract outline in (A) adapted from (Dean et al., 2009), the pre-leptotene cell was traced from (Endo et al., 2015), and all other cell images were adapted from (Larson et al., 2018).

To test for variation in evolutionary rates across spermatogenesis, we compared *d*_N_/*d*_S_ for genes associated with testis somatic cells, diploid spermatogonia, early meiosis (pre-leptotene), meiotic spermatocytes, postmeiotic spermatids, and elongating spermatids (Figure 3; Table 1) based on single-cell RNAseq data from house mice (Green et al., 2018). Genes enriched in testis somatic cells and early and middle spermatogenesis stages (spermatogonia, pre-leptotene, and spermatocytes) had median *d*_N_/*d*_S_ less than, or similar to, that of the genome-wide average, while genes enriched during the postmeiotic spermatid and elongating spermatid stages showed higher *d*_N_/*d*_S_ (Table 1; Figure 3A; FDR-corrected Wilcoxon rank-sum tests P < 0.05). These results suggest that more rapid protein sequence divergence in late spermatogenesis underlies rapid divergence of testis genes, consistent with results for both closely related mouse lineages (Good & Nachman, 2005; Kopania et al., 2022) and more broadly across mammals (Murat et al., 2022).

### Both positive selection and relaxed purifying selection contributed to the rapid divergence of some reproductive genes

The rapid evolution of genes enriched in seminal vesicles and during late spermatogenesis may result from relaxed purifying selection, positive directional selection, or both. To test for gene-wide positive selection across our murine phylogeny, we used the BUSTED model from HyPhy (Murrell et al., 2015). We ran BUSTED three times, once with no additional parameters, once with synonymous rate variation incorporated into the model, and once with both synonymous rate variation and multiple nucleotide substitutions incorporated into the model. We then calculated a model-averaged p-value, which weights p-values based on the likelihoods of different models (Lucaci et al., 2023). BUSTED, both with and without synonymous rate variation, found evidence for positive selection in over 40% of genes. However, when we added multiple nucleotide substitutions into the model, only 6% of genes showed evidence for positive selection (Supplemental Table S8). Ignoring multiple nucleotide substitutions can drastically increase false positive rates in tests for selection, and this problem may be particularly common when trees have short branches, such as in our dataset (Lucaci et al., 2023). Estimates of *d*_N_/*d*_S_ and p-values from the three separate BUSTED models are reported in Supplemental table S9. Here we discuss results based on the model-averaged p-values, because this approach tends to reduce false positive rates while providing similar statistical power to models with fewer parameters (Lucaci et al., 2023).

For the seminal vesicles, 14% of genes showed evidence for positive selection (Murrell et al., 2015; Lucaci et al., 2023), suggesting that positive selection may contribute to the rapid divergence of some seminal vesicle genes. However, this proportion of seminal vesicle genes under positive selection was not significantly different from the genome-wide average of 17% (FDR-corrected P > 0.05; Table 1; Supplemental Figure S3). Similarly, rapidly evolving round spermatids and elongating spermatids had some genes with evidence for positive selection, but did not have significantly different proportions of genes with evidence for positive selection than the genome-wide average (Figure 3B; Table 1; Pearson’s chi-squared FDR-corrected P > 0.05).

Many male reproductive genes are highly tissue-or cell-type-specific, leading to the prediction that they may be under relaxed purifying selection. We identified the proportion of cell-type-enriched genes that were testis-specific to estimate the degree to which relaxed constraint may shape molecular evolution across spermatogenesis stages. Early-stage diploid spermatogonia had proportionally fewer testis-specific genes, whereas midto late-stage spermato-genesis cell types had proportionally more testis-specific genes (Figure 3C; Table 1; FDR-corrected Pearson’s chisquared P < 0.05). Genes enriched in round spermatids had the highest proportion that were testis-specific (Figure 3C; Table 1). Positive selection and relaxed purifying selection are not mutually exclusive, and testis-specific genes may be freer to diverge rapidly under positive selection because they are not constrained by functions in other tissues (Murat et al., 2022). However, testis-specific genes had a similar proportion of genes under positive selection compared to the genome-wide average (testis-specific: 13%; genome-wide: 17%; FDR-corrected P > 0.05; Table 1). Thus, some genes enriched in reproductive tissues and cell types show evidence for positive selection, but rapidly evolving reproductive tissues and cell types do not show elevated rates of positive selection. Relaxed purifying selection due to tissue specificity may be a major contributor to the rapid divergence of late spermatogenesis genes.

### Spermatogenesis genes showed accelerated evolutionary rates in species with low levels of sperm competition due to relaxed purifying selection

We next used *RERconverge* (Kowalczyk et al., 2019) to test if changes in the intensity of sperm competition were associated with lineage-specific shifts in molecular evolution. Because small testes mass was the trait that showed convergent optimum shifts in our OU model, we first set small testes species as the foreground branches (Figure 1, orange and brown groups denoted by †). Of the 11,775 protein-coding genes included in our post-filtering dataset, 11,588 genes had enough total species and independent foreground lineages included in their alignments to run *RERconverge*. Of these 11,588 genes, five showed significantly accelerated evolutionary rates associated with lower testis mass after correcting for multiple tests, and all five had expression patterns or functional annotations indicative of a role in male reproduction (positive *RERconverge* Rho values; FDR-corrected P < 0.05; Table 2). We also looked for functional enrichment among the most accelerated genes based on *RERconverge* p-values. Although only five genes had p-values < 0.05 after multiple test correction, many genes that rank near the top of *RERconverge* analyses can still show a tendency to be evolving more rapidly in foreground lineages (Kowalczyk et al., 2020). Testing for functional enrichment among top ranking genes, even if they are not individually significant, can determine if accelerated evolutionary rates associated with small testes mass is a broader pattern among genes involved in reproductive processes or if this pattern is limited to just a few genes. The 10 most accelerated genes had known functions in reproduction or were highly expressed in the testes (Table 2). The top 250 most accelerated genes were significantly enriched for “male gamete generation”, “spermatogenesis”, and “gamete generation” gene ontology (GO) terms (Figure 4; Supplemental Table S10; FDR-corrected P < 0.01). To ensure that this result was statistically robust, we used a combination of phylogenetic permutations and simulations [i.e., permulations (Saputra et al., 2021)] and found that terms related to reproduction were still among the most highly enriched GO terms (Supplemental Table S10). These results suggest that accelerated evolutionary rates in lineages with small testes mass occur across many genes involved in male reproduction, and the five genes with significant *RERconverge* p-values likely reflect a broader pattern that is not unique to those genes.

**Table 2.** The top 10 most accelerated genes in small relative testes mass species based on *RERconverge*. Higher Rho values indicate a stronger positive correlation between relative molecular evolutionary rate and lineages with shifts to smaller relative testes mass. P.adj is the p-value from *RERconverge* corrected for multiple tests. Expression patterns are based on the Bgee suite (Bastian et al., 2021). Reproductive functions are based on Uniprot (The UniProt Consortium, 2020) and the Mouse Genome Informatics database (Baldarelli et al., 2024) unless noted otherwise.

| Gene symbol | Rho | P.adj | Expression Patterns | Reproductive Function |
| --- | --- | --- | --- | --- |
| *Zcchc13* | 0.3548 | 8.35E-03 | High in testis | Homozygous mutants show normal fertility |
| *Stk31* | 0.3386 | 8.35E-03 | Highest in spermatocyte; also highly expressed in other gonadal and zygotic tissues | Homozygous mutants show normal fertility |
| *Akap4* | 0.3171 | 3.98E-02 | Testis-specific (especially spermatid) | Part of sperm fibrous sheath structure; involved in sperm motility |
| *Sel1l2* | 0.3537 | 4.16E-02 | Highly expressed in spermatid, spermatocyte, and seminiferous tubules | Unknown |
| *Ppp2r1b* | 0.3129 | 4.16E-02 | Expressed in several developing tissues, including indifferent gonad; also expressed in seminal vesicle and spermatid | Essential for proper meiosis in spermatocytes; homozygous deletion mice infertile (54) |
| *Sox30* | 0.2837 | 6.30E-02 | High in testis | Activates postmeiotic genes involved in spermiogenesis |
| *Ddx43* | 0.2651 | 1.37E-01 | Highest in spermatocyte; also highly expressed in other gonadal and zygotic tissues | Chromatin remodeling; homozygous deletion mice infertile (55) |
| *Calr3* | 0.2706 | 1.70E-01 | Highest in spermatid; also highly expressed in other gonadal and zygotic tissues | Required for zona pellucida binding and fertilization |
| *Angel1* | 0.2658 | 1.70E-01 | Highest in spermatid; also highly expressed in other gonadal and zygotic tissues | Unknown |
| *Ybx2* | 0.2739 | 1.74E-01 | Highest in seminiferous tubule; also highly expressed in other gonadal and zygotic tissues | Required for late stages of spermatogenesis; required for both male and female fertility |

Because genes involved in reproduction tend to evolve rapidly, this enrichment for reproduction-related terms among genes accelerated in small testes species could be due to overall rapid divergence of these genes, rather than being specific to genes accelerated in small testes species. To test if this enrichment for spermatogenesis genes was unique to genes with accelerated evolutionary rates in small testes mass species, we ranked genes by their *RERconverge* correlation score and tested for enrichment of spermatogenesis genes among genes falling within sliding windows using a tricube moving average, which down-weights values at the ends of the window to prevent extreme values from having a disproportionate effect on the moving average (Kowalczyk et al., 2020). This showed that only genes accelerated in small testes species were enriched for the spermatogenesis GO term (Figure 4A).

**Figure 4.**
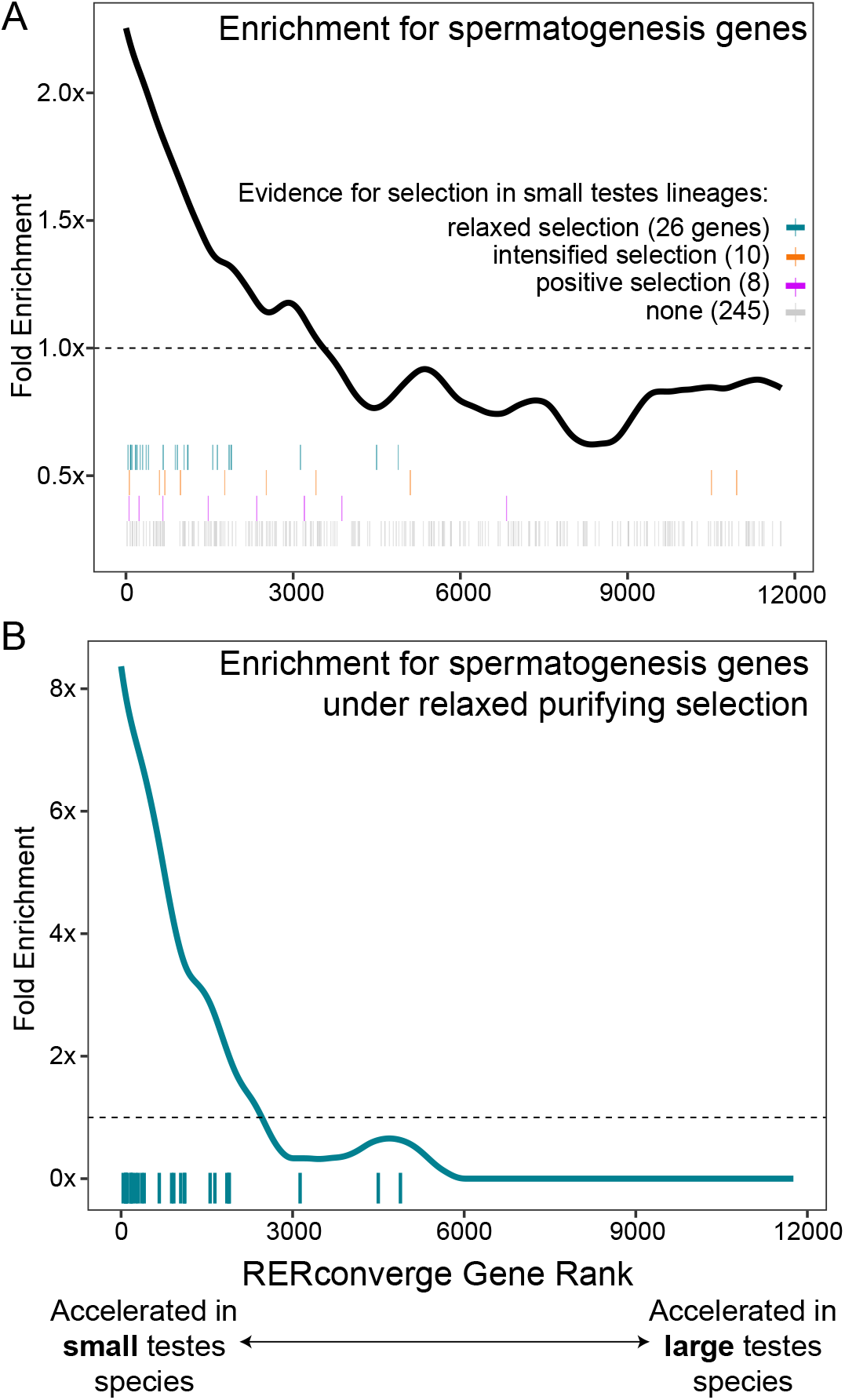
Enrichment for genes in the spermatogenesis GO category based on *RERconverge* rank. The x-axis indicates every gene included in the *RERconverge* analysis, ordered by their correlation score, with genes on the far left indicating those accelerated in small testes species. The line indicates enrichment for genes in the spermatogenesis GO category in windows along the x-axis based on a tricube moving average. The horizontal dashed line is at 1x and therefore represents no over-or under-enrichment. Each tick mark indicates a gene in the spermatogenesis GO category, colored based on evidence for selection, and the placement of these ticks along on the y-axis is arbitrary. (A) Enrichment for the spermatogenesis GO category in sliding windows across all genes included in the *RERconverge* analysis. (B) Enrichment for genes in the spermatogenesis GO category with evidence for relaxed selection [i.e., observed/expected numbers of genes represented by teal tick marks in (A) in sliding windows along the x-axis].

We also ran *RERconverge* with different sets of foreground species, including with large relative testes mass (highest 10%) species as foreground, and with relative testes mass as a continuous trait. For the continuous trait model, we also found that spermatogenesis genes and testis-specific genes were accelerated in smaller testes species, but not in larger testes species (Supplemental Figure S4). When we assigned foreground branches based on large testes mass species, neither the most accelerated genes nor the most decelerated genes were enriched for any GO terms (FDR-corrected P > 0.05). Overall, these comparisons suggest that relaxed purifying selection in low sperm competition species, rather than positive selection in species with high sperm competition, contributes more to the interspecific variation in evolutionary rates of spermatogenesis genes in murines.

To test this hypothesis, we tested for positive selection acting on spermatogenesis genes on branches leading to small testes species using HyPhy BUSTED-PH. We also tested for relaxed purifying selection in these lineages using HyPhy RELAX. We found evidence for relaxed purifying selection acting on 26 spermatogenesis genes in small testes species, and the majority of these were also accelerated in small testes species (Figure 4A, teal tick marks; Supplemental Table S11). We saw an even stronger enrichment for genes that were both under relaxed selection and in the spermatogenesis GO category among genes accelerated in small testes species (8-fold enrichment, Figure 4B), compared to the enrichment for genes in the spermatogenesis GO category regardless of selection mode (2-fold enrichment, Figure 4A). In contrast, only 8 spermatogenesis genes showed evidence for convergent episodic selection in small testes species, and these were more spread out across the range of *RERconverge* output values rather than concentrated among genes accelerated in small testes mass species (Figure 4A, red tick marks). RELAX also identifies genes under intensified selection in test branches; we only found 10 genes with evidence for intensified selection associated with small relative testes mass species (Figure 4A, purple tick marks).

## Discussion

Many of the most remarkable examples of rapid phenotypic and molecular evolution are related to male reproduction (Breed, 2004; Clark et al., 2006; Pitnick et al., 2009; Lüpold et al., 2016), but few studies have directly tested how interspecific differences in inferred post-mating sexual selective pressures act on both phenotypic and molecular evolution. Our dataset consisting of genomic and reproductive trait data from 78 species allowed us to address this in a rigorous phylogenetic framework. Below we first discuss insights into the evolution of sperm traits relative to inferred variation in mating system (i.e., intensity of sperm competition) across a large sample of murine rodents. We then consider how this phenotypic variation influences underlying patterns of protein-coding evolution across male reproductive tissues, providing a rare integration of molecular and phenotypic evolution across a large radiation of mammals.

### The evolution of male reproductive traits in murine rodents

We found evidence for the repeated evolution of smaller relative testes mass, suggesting that most murines have a high frequency of female multiple mating but with repeated shifts to mating systems with less frequent female remating (Figure 1). We also identified several sperm morphological traits associated with relative testes mass (Figure 2). Our findings suggest that sperm competition selects for the presence of an apical hook on the sperm head, as well as longer and more angled hooks, consistent with previous studies (Immler et al., 2007; Varea-Sánchez et al., 2016; Pahl et al., 2018). For example, compared to a previous study that sampled fewer murine species with less sperm diversity (Pahl et al., 2018), we found consistent support that relative testes mass is associated with hook morphology and sperm tail length, but in contrast we did not find an association with sperm head area.

Our data suggest that sperm hook evolution is relevant to sperm competition in murines, but hook function and its role in post-copulatory sexual selection remains a mystery. In some species, the hooks attach sperm from the same male to form sperm trains, which speed the transfer to the egg (Moore et al., 2002). This was thought to be the main function of the hook (Immler et al., 2007), but other works have revealed that many species rarely form sperm trains, and trains that do form disaggregate quickly and do not show increased swimming speed or fertilization success (Firman & Simmons, 2009; Firman et al., 2013; Tourmente et al., 2016). Alternatively, hooks may improve sperm swimming speed (Varea-Sánchez et al., 2016) or attach to the female reproductive tract to avoid being flushed out (Firman & Simmons, 2009). The hook also contains part of the acrosome, which contains enzymes that break down the zona pellucida egg coat layer, and therefore could facilitate sperm attachment to the egg cell membrane (Breed, 1984, 2004). Regardless of the hook’s proximal functions, its presence and morphology are likely to play an important role in sperm competitive ability (Hook et al., 2021; Teves & Roldan, 2022).

It was not feasible to directly measure levels of sperm competition for most species in our dataset, so we used relative testes mass as a proxy for the intensity of sperm competition. There is a positive correlation between the intensity of sperm competition and relative testes mass in many vertebrate groups (Harcourt et al., 1981; Birkhead et al., 1993; Ramm et al., 2005; Firman & Simmons, 2008), including some murines (Ramm et al., 2005; Firman & Simmons, 2008). However, this correlation is probably imperfect, because testes perform functions beyond sperm production, and changes in testes architecture can lead to increased sperm production without increased testes size (Pitnick & Markow, 1994; Ramm & Schärer, 2014). Furthermore, some large testes species display behaviors associated with monogamy, such as biparental care in *Mus spretus* (Cassaing et al., 2009). Relative testes mass also may not reflect other components of post-mating sexual selection that influence sperm morphology, such as cryptic female choice (Firman et al., 2017). For example, cryptic female choice may also act on sperm hook morphology, especially if the hook is involved in interactions with the female reproductive system (Firman & Simmons, 2009).

### The relationship between phenotypic and molecular evolution in response to sperm competition

We showed that the rapid molecular divergence of male reproduction in murine rodents primarily occurred in testis-specific genes, and genes enriched in the seminal vesicles and postmeiotic spermatogenesis cell types, consistent with several previous studies [Figure 3; Table 1; (Dean et al., 2009; Larson et al., 2016; Kopania et al., 2022; Murat et al., 2022)]. These genes may play important roles in both sperm competition and sexual conflict. For example, the seminal vesicles produce secretions involved in copulatory plug formation, which is important for fertilization success and to limit female remating (Dean et al., 2009; Mangels et al., 2016), and are also enriched for immune function genes (Dean et al., 2009), which evolve rapidly in primate ejaculates (Good et al., 2013) and in murine rodents (Roycroft et al., 2021a). Postmeiotic spermatogenesis genes may be subject to positive directional selection on diverse aspects of sperm head morphology [(Pitnick et al., 2009); Figure 2], which develops during the later stages of spermatogenesis. An important caveat is that our sets of tissue-specific or enriched genes came from house mouse (*M. musculus*) data, and therefore may not reflect the expression patterns of these genes in every species in our tree. Genes involved in reproduction, with more specific expression patterns, and with sex-biased expression tend to show more rapid turnover in their expression patterns (Brawand et al., 2011; Soumillon et al., 2013; Murat et al., 2022). However, many reproductive genes also show conserved expression patterns across mammalian spermatogenesis (Chalmel et al., 2007; Murat et al., 2022), and our filtering to only include genes with confident 1:1 orthologs between mouse and rat likely eliminated novel genes and genes with rapid gene family evolution that may be more likely to have divergent expression patterns. Therefore, we expect that our gene sets likely give a reasonable approximation of molecular evolutionary patterns across different tissue and cell types in murines.

Previous studies have shown strong evidence for positive selection acting on a subset of reproductive genes, which plays an important role in their overall rapid divergence (Swanson et al., 2001; Clark & Swanson, 2005; Ahmed-Braimah et al., 2017; Dean et al., 2017; Roycroft et al., 2021a), but relaxed purifying selection likely also plays an important role in the rapid divergence of reproductive genes (Dean et al., 2009; Larson et al., 2016; Schumacher & Herlyn, 2018; Kopania et al., 2022; Murat et al., 2022). We did not find significantly higher proportions of genes under positive selection in the rapidly evolving seminal vesicles or postmeiotic spermatids (Table 1), suggesting that relaxed purifying selection may play a more important role in their rapid divergence. However, relaxed purifying selection and positive selection are not mutually exclusive, and the relaxation of purifying selection may allow for the rapid fixation of mutations under positive selection (Larracuente et al., 2008). Additionally, selection can act on structural variation in addition to sequence variation, and copy number expansion may be under positive selection in some reproductive genes (Ramm et al., 2008a)(Podlaha et al., 2005; Nam et al., 2015). We focused on 1:1 orthologs to compare gene sequence divergence, but it is possible that some genes in our dataset were evolving under positive selection on structural rather than sequence variation.

We predicted that lineages with a higher intensity of sperm competition would show more rapid molecular evolution of reproductive genes that tend to evolve under positive directional selection, such as seminal fluid proteins and genes involved in fertilization. However, we observed a different pattern, with a subset of spermatogenesis and testis-specific genes tending to evolve more rapidly in small testes species due to relaxed purifying selection, but with no enrichment of any reproductive genes among those rapidly evolving in large testes species (Figure 4A; Supplemental Figure S4). It is notable that strong purifying selection, rather than positive selection, under high sperm competition appears to have the strongest influence on lineage-specific patterns of molecular evolution of genes involved in male reproduction. The ten most accelerated genes in small testes species are all highly expressed in house mouse testes (Bastian et al., 2021), and some have known essential functions in house mouse male fertility (Table 2). However, others do not affect fertility in mouse mutational studies or have no known functions in reproduction (e.g., Zcchc13, Stk31, Sel1l2, Angel1), and therefore provide candidates for future studies investigating variation male fertility especially in the context of sperm competition.

We showed that rapid divergence of reproductive genes in small testes species was largely due to relaxed purifying selection in these lineages (Figure 4B). Although sperm competition is typically assumed to be associated with positive selection, our results show that sperm competition also influences the intensity of purifying selection on many genes, and that shifts in these pressures can be a major driver of molecular evolution. This has been predicted theoretically (Dapper & Wade, 2016; Dapper & Wade, 2020), and recent studies have found similar results in primates (Ports & Jensen-Seaman, 2023; Bowman et al., 2024) and Drosophila (Patlar et al., 2021), suggesting that stronger purifying selection under sperm competition may be a consistent pattern across diverse taxa. Strong purifying selection on spermatogenesis genes in large testes species is also consistent with phenotypic observations. For example, there is a negative correlation between intraspecific variation in sperm morphology and relative testes mass in rodents and passerine birds, suggesting that sperm competition pressures may reduce variation in sperm morphology (Immler et al., 2008; Varea-Sánchez et al., 2016). In the small testes murine species *Notomys alexis* and *N. fuscus*, there is high intraspecific variation in sperm hook morphology, with some having small hooks or ventral processes and others lacking these structures altogether (Breed et al., 2020a). Thus, sperm competition may impose purifying selection on genes involved in sperm development to facilitate a higher proportion of high-quality sperm in the ejaculate. We suggest that this represents a more complete understanding of the molecular evolution of reproductive genes, in which a small subset of genes evolve rapidly due to positive directional selection across most taxa, while many others are highly conserved in lineages with intense sperm competition due to strong purifying selection but evolve rapidly in lineages where this purifying selection is relaxed.

## Data availability

Exome data and assemblies are available at accession PRJNA1109543. Raw sequence reads for all samples generated by Bioplatforms Australia are available from BPA Data Portal as part of the Oz Mammals Genomics Initiative using Library IDs provided in Supplemental Table S1 (https://data.bioplatforms.com). Scripts for assembly, alignment, and phylogenetic inference are available at https://github.com/goodest-goodlab/murinae-seq/tree/master. Scripts for all other analyses, the pruned 78 species tree, and lists of genes associated with reproductive tissues and cell types are available at https://github.com/ekopania/murine-sperm-analyses/tree/main.

## Author Contributions

J.A.E., K.C.R., and J.M.G. conceived the project. A.S.A., J.A.E., and K.C.R. acquired specimens. S.M.E.M., C.M.C., and E.J.R. collected the genomic data. W.G.B. and E.E.K.K. compiled the phenotypic data. G.W.C.T. and C.R.H. performed the assembly, alignment, and phylogenetic inference. E.E.K.K. analyzed the data with support from N.L.C. N.L.C., J.A.E., K.C.R., and J.M.G. funded the projec E.E.K.K. and J.M.G. wrote the manuscript with input from all authors.

## Acknowledgements

We thank the Good and Clark labs, Doug Emlen, Lila Fishman, Erica Larson, Travis Wheeler, and two anonymous reviewers for helpful feedback on this study. We thank Ana Paula A Assis for assistance with the OU models. We thank the Museum of Vertebrate Zoology, Field Museum of Natural History, Louisiana State University Museum of Natural Science, Western Australia Museum, South Australia Museum, University of Kansas Biodiversity Research Center, Australian Biological Tissue Collection, Museum and Art Gallery of the Northern Territory, University of Canberra, Queensland Museum, Muséum National d’Histoire Naturelle, and Museums Victoria for providing tissue samples.

