## Supplemental Material for "Sperm competition intensity shapes divergence in both sperm morphology and reproductive genes across murine rodents"

**Supplemental Methods**

**DNA extraction**

To account for ethanol preservation, we made the following modifications to the Qiagen DNeasy Kit. We rehydrated tissues with two 30-minute incubations of 1mL 1× STE buffer followed by three to five minutes of vortexing (Bi et al., 2013). To lyse samples, we added 20µL 1M DTT and 10µL 0.5M EDTA (Shapiro & Hofreiter, 2012), in addition to the Buffer ATL and proteinase K.

**Reproductive phenotype data collection**

Relative testes mass was reported as percent of body mass (paired testes mass / body mass; (Breed & Taylor, 2000; McLennan et al., 2017; Peirce et al., 2018; Breed et al., 2020). In some cases, only one testis was weighed and its mass was doubled to approximate paired testes mass (Pahl et al., 2018; Breed et al., 2019). We used percent of body mass rather than the observed/expected mass based on allometric studies to be consistent with previous studies in murines that reported relative testes mass as a percent of body mass (Breed & Taylor, 2000; McLennan et al., 2017; Pahl et al., 2018). The range of body mass values across murines is from about 10g to 1000g (Breed & Taylor, 2000), so the negative allometry of testes mass with body mass likely has a small effect at this scale (Kenagy & Trombulak, 1986). Sperm morphological traits were measured from scanning electron microscope images (McLennan et al., 2017; Pahl et al., 2018; Peirce et al., 2018; Breed et al., 2019). Sperm head length was measured from the base of the head to the base of the apical hook, and head width was measured perpendicular to head length at the widest part of the head (McLennan et al., 2017; Breed et al., 2019). Sperm head area was measured by tracing the outer surface of the sperm head including apical hooks and ventral processes and using an area tool (Pahl et al., 2018). Apical hook and ventral processes lengths were measured from the base to the tip of the hook along the concave surface of the hook (McLennan et al., 2017; Pahl et al., 2018; Breed et al., 2019). For sperm with multiple ventral processes, the length of the longest ventral process was used (McLennan et al., 2017). Apical hook and ventral processes angles were reported as the angle between the tangent line from the tip to the base of the hook and the line along the sperm head longitudinal axis (Immler et al., 2007; McLennan et al., 2017; Pahl et al., 2018). More details can be found in the original references (Supplemental Table S2).

**Models for phylogenetic analyses**

We used three different correlation structures: Brownian motion, Pagel’s ƛ (Pagel, 1999), and an OU model (Martins & Hansen, 1997). We defined these correlation structures for our dataset using the functions *corBrownian*, *corPagel*, and *corMartins* with starting values set to 1. Under the Brownian motion model, variation in a trait is largely determined by phylogenetic history (Felsenstein, 1985), whereas the Pagel model allows traits to evolve independently of phylogeny (Freckleton et al., 2002). The Martins model represents an OU model, where evolution of a trait is somewhat determined by phylogenetic history but with adaptive peaks across the phylogeny (Martins & Hansen, 1997).

**Reproductive gene sets**

We tested if molecular evolutionary rates varied across male reproductive tissues using supplementary tables 1 and 2 from (Dean et al., 2009) to obtain lists of genes coding for proteins detected in each tissue (at least two peptides detected by mass spectrometry). Testes were not included in this proteomics dataset and also express most of the genes in the mammalian genome, so we focused on molecular evolutionary patterns of testis-specific genes from (Chalmel et al., 2007). We also tested if molecular evolutionary rates varied across spermatogenesis cell types by comparing *d*_N_/*d*_S_ for genes enriched for expression in different testes cell types. We used supplemental table S3 from (Green et al., 2018) to identify marker genes associated with each cell type cluster in their *Mus musculus* single-cell RNAseq dataset. We then used Figure 2B from (Green et al., 2018) to merge genes from these clusters into five cell type categories: spermatogonia, pre-leptotene, spermatocytes, spermatids, and elongating spermatids. We identified somatic cell marker genes using supplemental table S4D from (Green et al., 2018). Note that these datasets represent proteomics and expression patterns in *Mus musculus* only, because it was not feasible to generate proteomics or expression data for all species in our dataset. From these lists of genes associated with reproductive tissues or cell types, 1119 genes were excluded because they did not pass filtering, primarily due to lack of confident orthology between the mouse and rat references (Supplemental Table S3).

**Supplemental Figures**

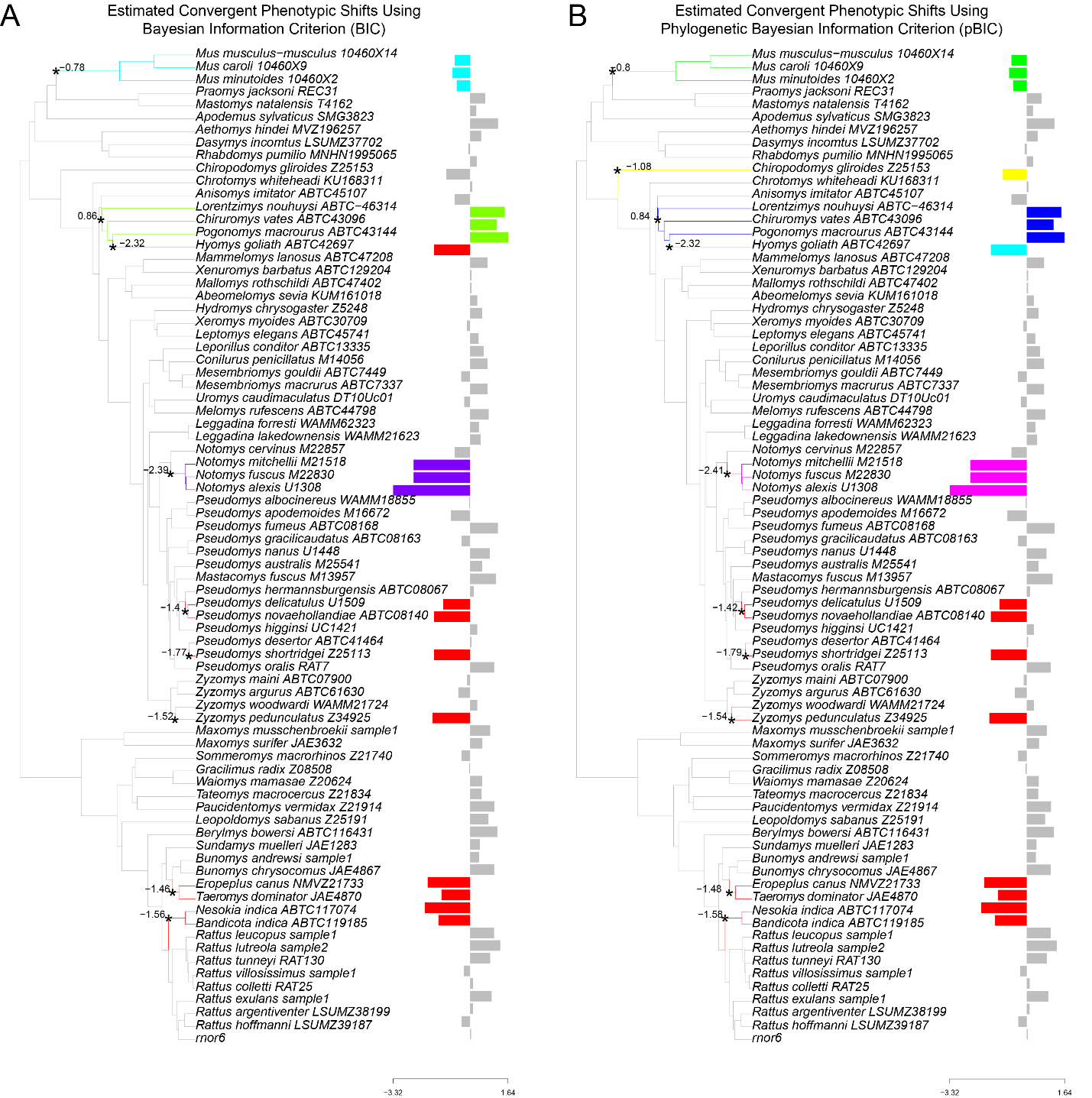
 **Figure S1.** Convergent shifts in relative testes mass phenotypic optima using different model selection tests in *l1ou*. Bar plots indicate normalized values of relative testes mass, with different colored bars indicating different adaptive peaks based on an OU model. Colored bars facing left indicate shifts to smaller testes and colored bars facing right indicate shifts to larger testes. Bars that are the same color represent convergent shifts to similar phenotypic optima based on the *estimate_convergent­_regimes* function in *l1ou*. Asterisks indicate points on the tree corresponding to these shifts in phenotypic optima, and values at asterisks indicate optimum trait values at these nodes. (A) Convergent regimes based on a Bayesian Information Criterion (BIC) test. (B) Convergent regimes based on a phylogenetic Bayesian Information Criterion (pBIC) test.

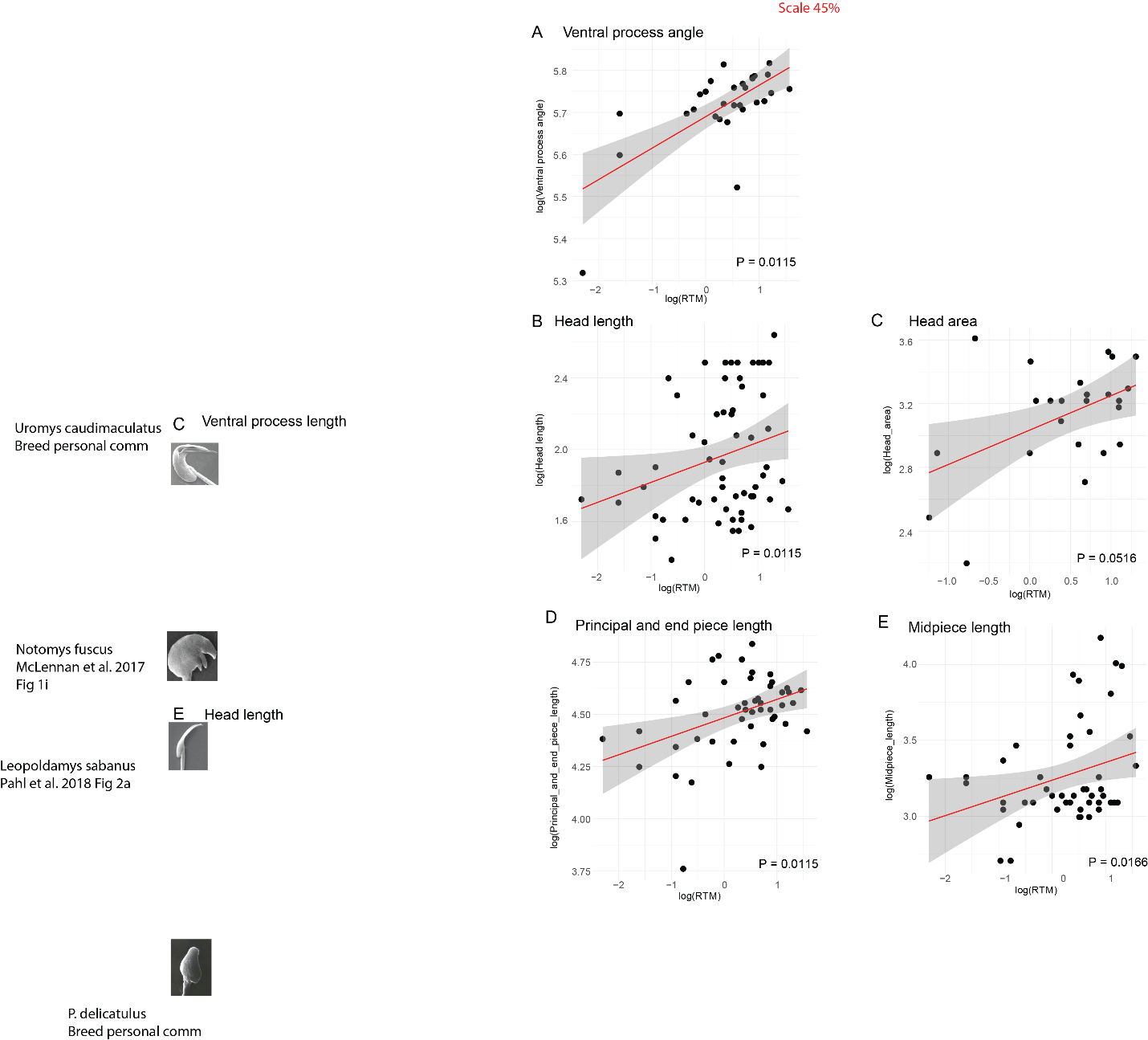

**Figure S2.** Correlations between relative testes mass (RTM) and sperm morphology traits. Red lines and gray areas show regressions and confidence intervals based on generalized linear models. P-values are based on phylogenetic generalized least squares analyses using a Brownian motion model with Benjamini-Hochberg correction for multiple tests. All lengths and widths were measured in µm, angles were measured in degrees, and relative testes mass was measured as a percent of body mass.

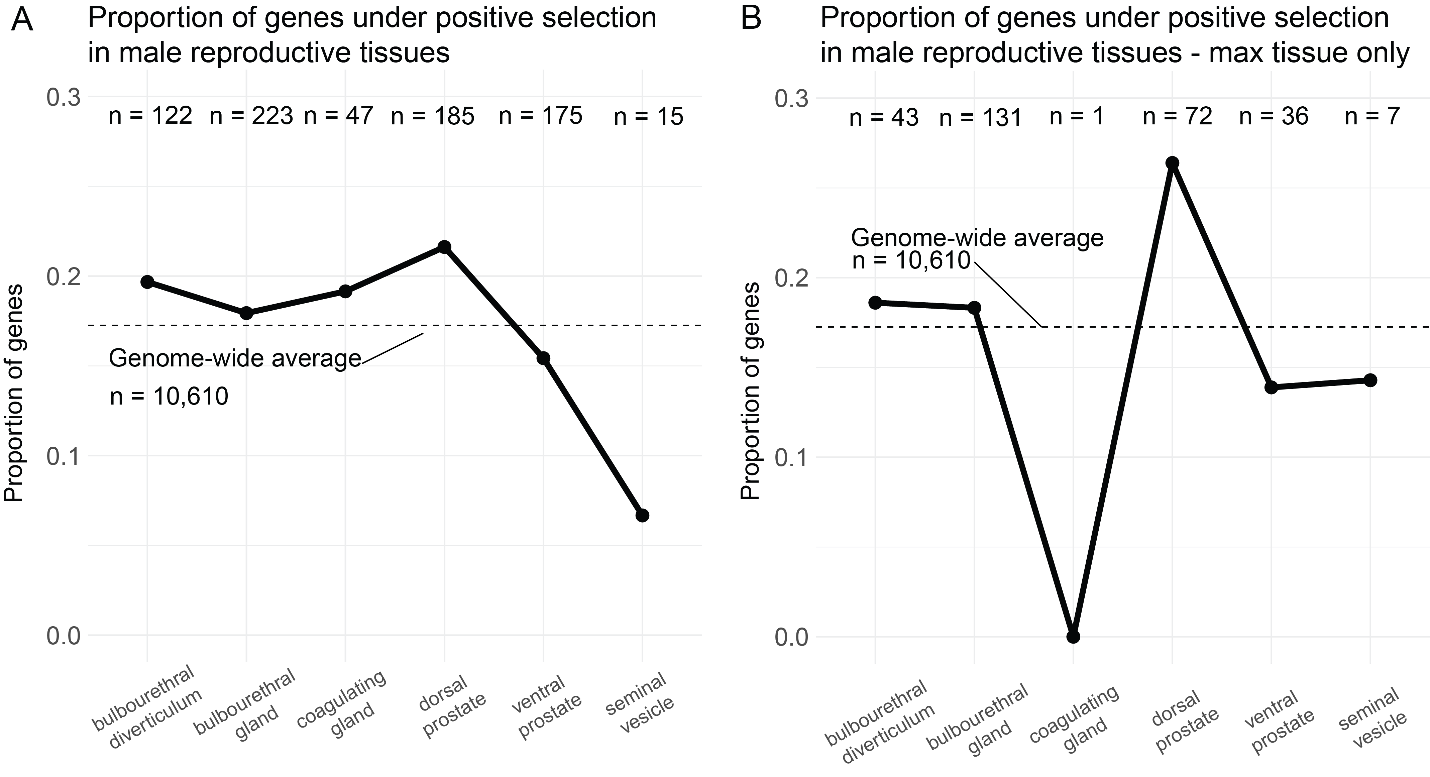

**Figure S3.** Proportion of genes enriched in reproductive tissues with evidence for positive directional selection. The numbers indicate the numbers of genes that were tested for positive selection in each cell type. For (A), genes were assigned to any tissues their protein products were detected in based on mass spectrometry data from Dean et al. 2009. For (B), genes were assigned to the tissue in which they were detected at the highest level based on mass spectrometry data from Dean et al. 2009 (i.e., a gene can be assigned to multiple tissues in A but can only be assigned to one tissue in B). No tissue was significantly different from the genome-wide average (FDR-corrected Pearson’s chi-squared P > 0.05).

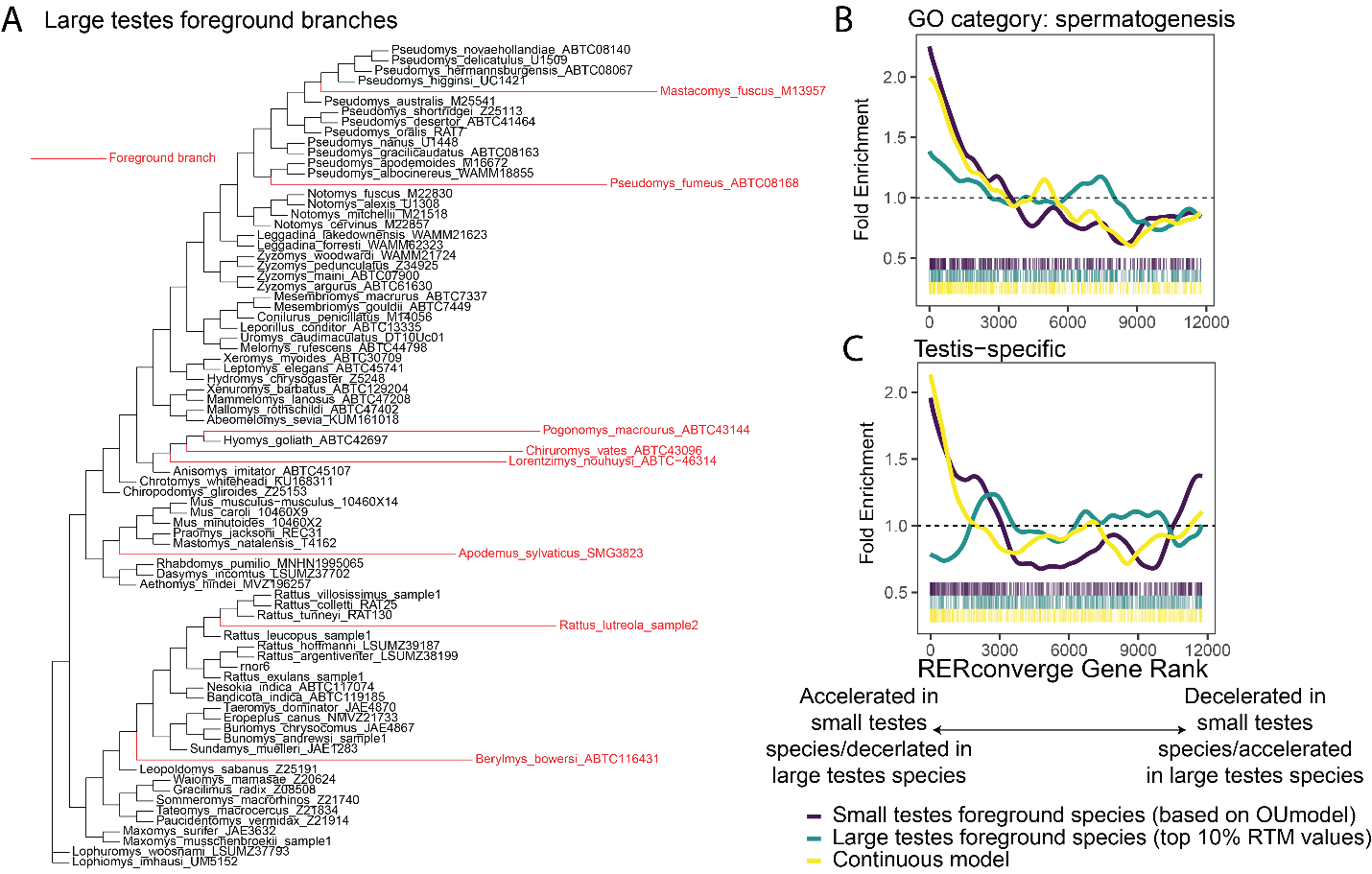

**Figure S4.** Comparison of *RERconverge* models. In addition to assigning foreground branches based on small testes species (main text), we also assigned foreground branches based on large testes species (A) and based on modeling relative testes mass as a continuous trait. (B) shows an enrichment for genes in the spermatogenesis gene ontology category for genes accelerated in small testes species in both the small testes foreground and continuous models. (C) shows an enrichment for testis-specific genes for genes accelerated in small testes species in both the small testes foreground and continuous models.
